# Ketone-loading as A Novel Sports Nutrition Strategy: Chronic Ketone Supplementation Elicits Further Favorable Metabolic Changes in Keto-adapted Mice

**DOI:** 10.1101/2023.04.30.537587

**Authors:** Sihui Ma, Shiori Onogi, Huijuan Jia, Hisanori Kato, Katsuhiko Suzuki

## Abstract

**Background:** A ketogenic diet (KD) induces nutritional ketosis (NS), benefits fatty acid oxidation (FAO), and favors moderate-intensity exercise capacity. The status that the body accommodates to produce and utilize ketone bodies (KB) and fatty acids as primary fuel sources is termed keto-adaptation. However, keto-adaptation requires time, while long-term KD also involves unfavored adverse effects. Exogenous ketone body (EKBs) administration has been introduced to elicit the advantages of NS. However, the direct use of EKBs fails to bring desired outcomes. We hypothesized that EKBs might only be effective during keto-adaptation.

**Methods:** Male C57BL/6J mice (n = 24) were divided into three groups: a control diet (Con, n = 8), a ketogenic diet (KD, n = 8), and a KD plus a ketone body (DL-β-Hydroxybutyric acid sodium salt, BHB) administration (KD+BHB, n = 8). After six weeks of KD administration, mice in the KD+BHB group receive BHB added into water bottles for another six weeks. Blood KB concentration is monitored throughout the experiment, while liver, gastrocnemius, and soleus mRNA are analyzed using RT-PCR.

**Results:** Both KD and KD+BHB induced and sustained NS and enhanced hepatic and muscular key genes regulating FAO. In addition, BHB administration upon keto-adaptation further increased circulating KB concentration and enhanced expressional levels of FAO-mediating genes (*ACO, HADH, ACADM*, and *MLYCD* in the gastrocnemius muscle; *ACO, HADH*, and *MLYCD* in the soleus muscle), and energy-regulating genes (*PPARA* and *PPARG*) in the liver and skeletal muscle compared to a KD.

**Conclusion:** Compared to KD alone, chronic administration of KBs upon keto-adaptation increased the expression of key genes that favor FAO or maintain energy homeostasis in the liver and skeletal muscle. Instead of directly using EKBs in non-keto-adapted individuals, it is encouraged to use EKBs upon keto-adaptation status to elicit their energy-utilizing effects.

**Highlights:** This is the first report to evaluate the metabolic effects using exogenous ketone bodies on keto-adapted individuals.

Administration of exogenous ketone body upon keto-adaptation furtherly increased circulating ketone bodies.

Administration of exogenous ketone body upon keto-adaptation individuals enhanced expression of genes related to fatty acid oxidation and energy hemostasis.

## 1. Introduction

A ketogenic diet (KD) induces a unique metabolic status named nutritional ketosis (NK), which is characterized by increased concentrations of ketone bodies (KBs) in the circulation (hyperketonemia) [1]. Previous studies report that KD benefits lipid oxidation through up-regulation of the expressional levels of rate-limiting enzymes in fatty acid mobilization and oxidation (FAO) [2],[3]. Therefore, strategies have been suggested to utilize KD to improve FAO, thus improving energy metabolism and exercise performance [4]. However, it takes time for the body to adapt to a KD with severe carbohydrate restriction measures, while long-term KD brings risk factors to the glucose metabolic system and the cardiovascular system, including glucose intolerance [5], hepatic steatosis, and hepatocyte apoptosis [6]. As a substitution, exogenous KB supplementation (EKBs) have been employed to induce NS in a few minutes without triggering ketogenesis by carbohydrate restriction, therefore reducing the difficulty and lowering the adverse effects [7]. However, attempts have been futile in employing exogenous ketone supplementations to directly benefits metabolism and performance, though nutritional ketosis is achieved in those [8],[9].

NS induced by KD or EKBs stand for two distinct metabolic statuses. Prolonged KDs induce multi-organ adaptations to utilize lipids and ketones as the primary energy sources, acclimating the body to a status termed keto-adaptation [10]. However, acute NS induced upon EKBs ingestion leaves the body remaining non-keto-adapted. Namely, the body is not adapted to change fuel preferences from glucose, therefore acute EKBs ingestion fails to elicit positive results. However, this hypothesis is yet validated. In the present study, we aim to inspect the effects of a six-week chronic EKBs in keto-adapted mice, to discuss the effects of EKBs upon keto-adaptation.

## 2. Materials and Methods

### 2.1 Animals and Diet

Male C57BL/6J mice (eight weeks old, total n = 24) were purchased from Takasugi Experimental Animals (Kasukabe, Japan). After one week of acclimation, mice were randomly divided into three groups: a control diet (Con, n = 8), a ketogenic diet (KD, n = 8), and a KD plus ketone body administration (KD+BHB, n = 8). Mice were provided a chow diet (Con; 10% (% kcal) protein, 80% CHO, 10% fat 3.8k/g, D19082304) or a KD (KD and KD+BHB, 10% protein, 0% CHO, 90% fat, 6.7kcal/g, D10070801, Research Diets Inc., NJ, USA). The fat source is cocoa butter. The experimental procedures were approved and followed the Guiding Principles for the Care and Use of Animals in the Academic Research Ethical Review Committee of Waseda University (2021-A13). Confounders are not controlled due to the small sample size. Experimenters could not be blinded due to the difference in fodder color and drink properties.

### 2.2 Exogenous Ketone Body Supplementation

DL-β-Hydroxybutyric acid sodium salt was used as EKB (H6501, Sigma-Aldrich, MA, USA). In the KD+BHB group, BHB was added to water bottles attempting to deliver ∼0.25 g/day for the first week as a “loading phase”, then 0.063 g/day as a “maintenance phase” for the remaining five weeks. This dosing schedule was designed to deliver a 60kg human-equivalent dose of 40 g/day dose during the loading phase and a 10 g/day dose during the maintenance phase per the body surface area mouse-to-human conversions using Meeh’s formula (11). Blood KB concentrations were monitored throughout the experiment using the FreeStyle Precision Neo system (Abbott, TX, USA). No exclusions were set.

### 2.3 RNA isolation and Real-time PCR

Total RNA was extracted from the gastrocnemius and soleus muscle using the RNeasy Fibrous Mini Kit, and from the liver using the RNeasy Mini Kit (Qiagen, CA, USA). Total RNA was reverse transcribed to cDNA using the High-Capacity cDNA Reverse Transcription Kit (Applied Biosystem (AB), CA, USA). PCR was performed with the Fast 7500 real-time PCR system (AB) using the Fast SYBR^®^ (AB) Green PCR Master Mix (AB). The thermal profiles consisted of 10 min at 95°C for denaturation followed by 40 cycles of 95°C for 3 s and annealing at 60°C for 15 s. 18s ribosomal RNA was used as the housekeeping gene, and the *ΔΔ*CT method was used. All data are represented relative to their expression as fold change to the Con group. Primer sequences are provided as supplementary data. No exclusions were set.

### 2.4 Statistical analysis

Data are presented as means ± standard deviation (SD). A one-way analysis of variance (ANOVA) was performed. Statistical analysis was done using Graphpad 9.0 (Graphpad, Ltd., CA, USA). Statistical significance was accepted as *p* < 0.05.

## 3. Results

### 3.1 Effects of BHB administration upon keto-adaption on circulating ketone body concentration, weight, and calorie intake

By the end of the study, blood KB concentrations were Con: 0.3±0.1, KD: 2.4±0.6, and KD+BHB: 3.3±0.8 mM. Significant differences are found between Con and KD, Con and KD+BHB, KD and KD+BHB. No significant differences were found among the groups on weight and calorie intake.

### 3.2 Effects of BHB administration upon keto-adaption in the liver

As shown in Figure 1A, both KD and KD+BHB enhanced the expressional levels of rate-limiting enzymes in lipid utilization. However, only KD+BHB increased the expression of *PPARA* and *PPARG* (Figure 1B) significantly compared to Con.

**Figure 1.**
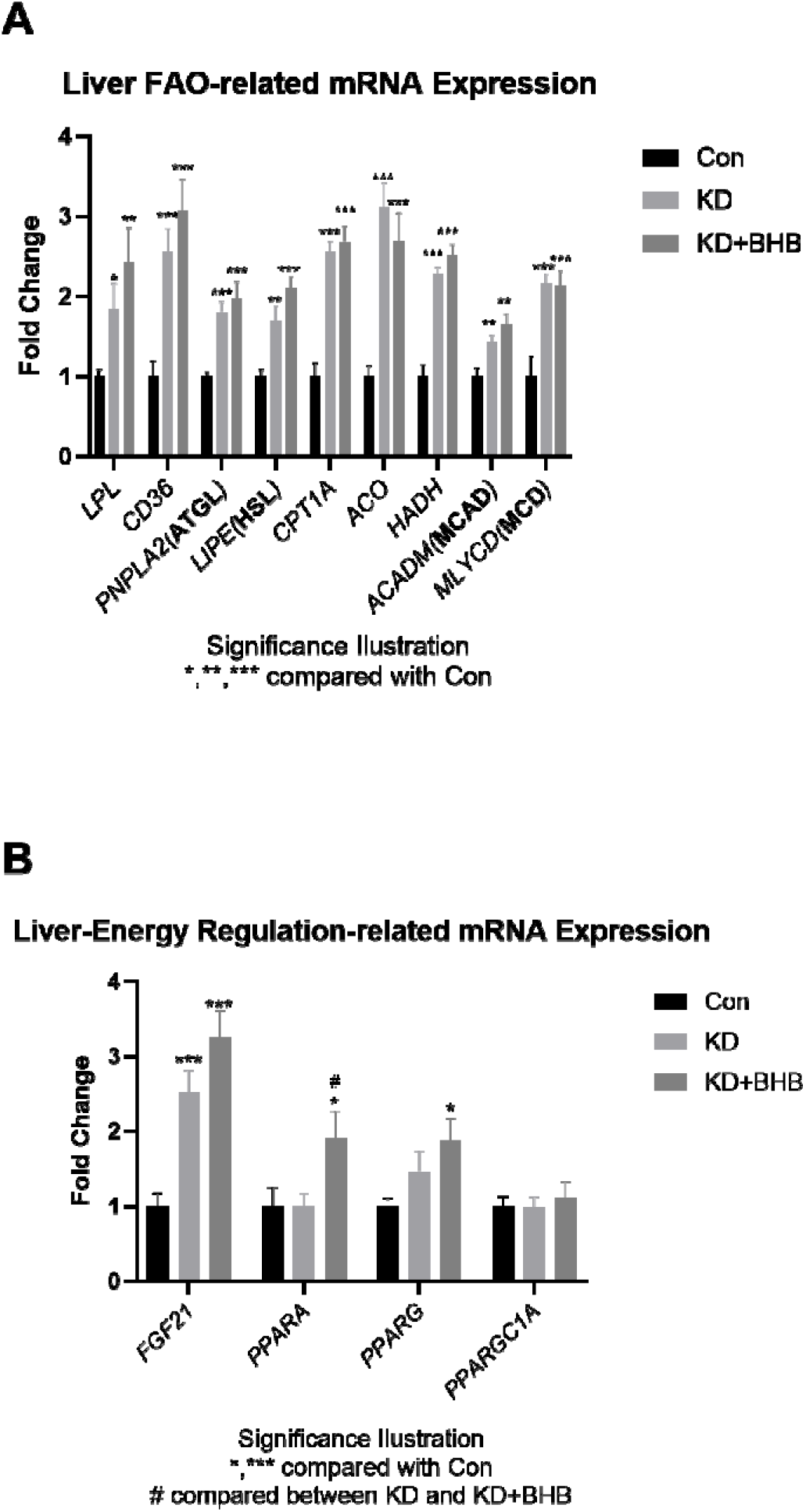
The effects of KD and KD+BHB administration in the liver. (A) KD and KD+BHB increased FAO-related mRNA expression. (B) Only KD+BHB increased *PPARA* and *PPARG* mRNA expression. N=8. *\*p*<0.05, *\*\*p*<0.01, and *\*\*\*p*<0.001, compared with Control, *#p*<0.05 compared with KD. *LPL*, lipoprotein lipase. *CD36*, cluster of differentiation 36. *PNPLA2(ATGL)*, adipose triglyceride lipase. *LIPE(HSL)*, hormone-sensitive lipase. *CPT1A*, carnitine palmitoyl transferase 1A. *ACO*, acetyl-CoA oxidase. *HADH*, hydroxyalkyl-CoA dehydrogenase. *ACAD(MCAD)*, medium-chain acyl-CoA dehydrogenase. *MLYCD(MCD)*, malonyl-CoA decarboxylase. *FGF21*, fibroblast growth factor 21. *PPARA*, peroxisome proliferator-activated receptor-alpha. *PPARG*, peroxisome proliferator-activated receptor-gamma. *PPARGC1A*, peroxisome proliferator-activated receptor gamma coactivator 1-alpha.

### 3.3 Effects of BHB administration upon keto-adaption in muscle tissues

As shown in Figure 2A, both KD and KD+BHB enhanced several rate-limiting enzymes of fatty acid utilization. Among them, *CD36, HADH*, and *MLYCD* expression in the fast-twitch muscle fiber gastrocnemius, *CD36, ACO, HADH, ACADM*, and *MLYCD* expression in the slow-twitch muscle fiber soleus, are both upregulated by a KD or KD+BHB, compared to Con. Moreover, KD+BHB further enhanced the expression levels of *ACO, HADH, ACADM*, and *MLYCD* in the gastrocnemius and enhanced the expressions of *ACO, HADH*, and *MLYCD* in the soleus compared to a KD. Furthermore, in the soleus muscle, only KD+BHB enhanced the expression of *CPT1A* significantly compared to Con.

**Figure 2.**
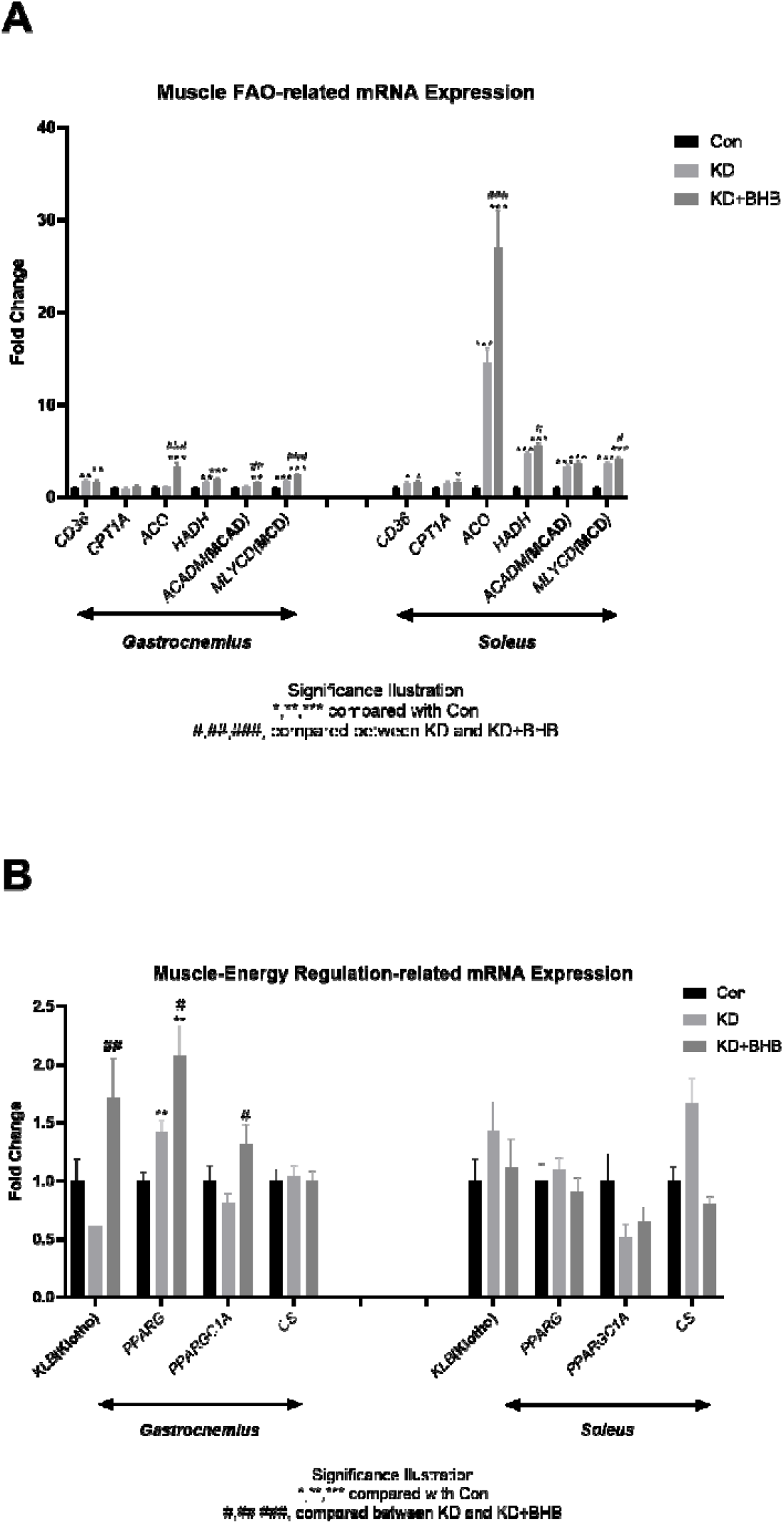
The effects of KD and KD+BHB administration in the muscle tissues. (A) KD and KD+BHB increased FAO-related mRNA expression. (B) Only KD+BHB increased *Klotho* and *PPARGC1A* expression. N=8. *\*p*<0.05, *\*\*p*<0.01, and *\*\*\*p<0*.*001*, compared with Control, *#p*<0.05, *##p*<0.01 and *###p*<0.001 compared with KD. *CD36*, cluster of differentiation 36. *CPT1A*, carnitine palmitoyl transferase 1A. *ACO*, acetyl-CoA oxidase. *HADH*, hydroxyalkyl-CoA dehydrogenase. *ACAD(MCAD)*, medium-chain acyl-CoA dehydrogenase. *MLYCD(MCD)*, malonyl-CoA decarboxylase. *KLB*, beta-Klotho. *PPARG*, peroxisome proliferator-activated receptor-gamma. *PPARGC1A*, peroxisome proliferator-activated receptor gamma coactivator 1-alpha. *CS*. citrate synthase.

As shown in Figure 2B, both KD and KD+BHB enhanced *PPARG* expression in the gastrocnemius compared to Con, while the enhancing effect of KD+BHB is more robust than KD’s. Moreover, only KD+BHB enhanced the expression of *Klotho* and *PPARA* in the gastrocnemius significantly compared to Con.

## 4. Discussion

Based on the theory that NS favors lipid oxidation, hypotheses have been made that those EKBs may favor overall metabolism, including enhancing exercise performance, without compromising carbohydrate reserves. However, most studies report disappointing results, including failure to improve exercise performance or impair exercise capacity, though FAO capacity is enhanced [8],[9]. Instead of acute KB administration to non-keto-adapted individuals, in the present study, we administered KBs to animals that are already keto-adapted. Our primary outcomes showed that KBs administration upon keto-adaptation further increased blood KBs concentrations compared to a KD. As secondary outcomes, we found that 1) hepatic and muscular genes involved with energy regulation were upregulated by KD+BHB, compared with KD, and 2) muscular genes involved with FAO were furtherly upregulated by KD+BHB, compared with KD.

PPARα regulates the expression of genes critical for lipid and lipoprotein metabolism and predominantly regulates fatty acid oxidation pathways, while PPARγ is a major activator of adipogenesis and fatty acid storage. Activation of hepatic PPARα and/or PPARγ favor overall lipid biosynthesis and catabolism, improving energy homeostasis [12]. Activation of muscular PPARγ benefits overall metabolism by inducing muscle-derived adiponectin, preventing insulin resistance, *etc*. PGC1α is a co-activator of PPARs and regulates mitochondrial biosynthesis [13]. In the present study, we found that the administration of KBs on keto-adapted animals increased the expression of PGC1α, indicating that this physiological phenomenon may favor mitochondrial function.

Klotho is one receptor of the energy-sensing molecule FGF21, and overexpression of Klotho has multiple preventive effects, including anti-aging, suppressing apoptosis, *etc*. [14]. KB administration upon keto-adaptation may elicit the advantages though inducing *Klotho* overexpression.

In our previous studies, a KD improves exercise capacity by enhancing FAO by upregulating expressional levels of rate-limiting enzymes in FAO, including *ACO, HADH*, and *MCD*. (2,3). In the present study, the above genes are further increased in BHB administration, indicating that KB administration upon keto-adaptation may additionally favor the exercise capacity.

Besides their role as an energy source, KBs also have signaling functions, linking environmental cues, such as starvation, to the body’s energy homeostasis [15]. KBs concentration in the KD+BHB group increased compared to KD, introducing a unique metabolic status where FAO capacity is further enhanced and may favor exercise performance.

Limitations exist. Though FAO capacity is enhanced in KD+BHB mice, we did not investigate the exercise capacity of the animals. Further study is encouraged to utilize the current results for applicational and clinical use.

## 5. Conclusion

This is the first report to use EKBs on keto-adapted animals. We found that chronic administration of KBs increased the expression of key regulators of lipid metabolism in the liver, and upregulated FAO-related mRNAs in skeletal muscle. Using EKBs upon keto-adaptation status is encouraged to elicit their energy-utilizing effects.

## Author contributions

Conceptualization, investigation, formal analysis, and writing - original draft: S.M. and S.O. Writing - review & editing: S.M, K.S, and H.J. Supervision: K.S.

